# Reduced inflammatory response and promoted multiciliated cell differentiation in mice protected by defective interfering influenza virus

**DOI:** 10.1101/2022.01.25.477719

**Authors:** Chang Wang, Rebekah Honce, Mirella Salvatore, Jianjun Yang, Nicholas M. Twells, Lara K. Mahal, Stacey Schultz-Cherry, Elodie Ghedin

**Affiliations:** Center for Genomics and Systems Biology, Department of Biology, New York University, New York, NY; Department of Infectious Diseases, St. Jude Children’s Research Hospital, Memphis, TN; Integrated Program in Biomedical Sciences, Department of Microbiology, Immunology, and Biochemistry, University of Tennessee Health Science Center, Memphis, TN; Department of Medicine, Weill Cornell Medical College, New York, NY; Department of Population Health Sciences, Weill Cornell Medical College, New York, NY; Department of Chemistry, University of Alberta, Edmonton, AB, CANADA; Systems Genomics Section, Laboratory of Parasitic Diseases, NIAID, National Institutes of Health, Bethesda, MD

## Abstract

Influenza defective interfering (DI) viruses have long been considered promising antiviral candidates because of their ability to interfere with replication-competent viruses and to induce antiviral immunity. However, the mechanisms underlying DI-mediated antiviral immunity have not been extensively explored. Here, we demonstrated interferon (IFN) independent protection conferred by influenza DI virus against homologous virus infection in mice deficient in type I and III IFN signaling. By integrating transcriptional and post-transcriptional regulatory data we identified unique host signatures in response to DI co-infection. DI-treated mice exhibited reduced viral transcription, less intense inflammatory and innate immune responses, and primed multiciliated cell differentiation in their lungs at an early stage of infection, even in the absence of type I or III IFNs. Overall, our study provides mechanistic insight into the protection mediated by DIs, implying a unifying theme involving inflammation and multiciliogenesis in maintaining respiratory homeostasis, and reveals their IFN-independent antiviral activity.

## INTRODUCTION

Defective interfering (DI) virus particles carry defective viral genomes (DVGs) but normal structural proteins and can interfere with the propagation of homologous replication-competent viruses [1-3]. Although different forms of DVGs exist among RNA viruses, DVGs in the influenza virus—a negative-sense RNA virus with a segmented genome—typically contain internal deletions within the genomic segments while preserving the 5’ and 3’ termini [4-8]. Naturally occurring DI viruses have been observed during virus propagation [1, 2, 9-12] and in human infections [13-15], and are believed to play an important role in modulating infection outcome [14-18]. DIs presumably compete with standard virus for components essential for replication, but also possess the ability to induce antiviral immunity and facilitate the establishment of persistent infections (reviewed in [5, 6, 19]). The immunostimulatory activity of DI viruses resides in their strong pathogen-associated molecular patterns (PAMPs), which can be recognized by pattern recognition receptors (PPRs) and subsequently induce interferon (IFN) production [15, 20, 21]. Additional sequence features, as demonstrated in copy-back DVGs, can also promote the binding and activation of PPRs [22].

IFNs are an important group of innate antiviral cytokines induced in response to viral infections. Among three IFN families identified so far, type I and III IFNs play a prominent role in defending the respiratory epithelial barrier [23]. Despite their distinct receptors, both type I and III IFNs signal through the JAK/STAT pathway to promote the transcription of a large spectrum of interferon-stimulated genes (ISGs) that exert various activities to combat viral infection (reviewed in [24-27]). Although substantially overlapping properties of type I and III IFNs renders them seemingly redundant [23], recent studies have revealed spatial and kinetic differences in their signaling and induction in the respiratory tract, highlighting their distinct features [28-33].

The immunostimulatory ability of DI viruses and the importance of IFNs in the innate antiviral response have motivated studies to explore the role the type I IFN response plays in DI-mediated antiviral protection *in vivo* [34, 35]. A study of mice lacking the type I IFN receptor challenged concurrently with an influenza A virus (IAV)-derived DI virus (i.e., DI244) and a wild-type (WT) IAV suggests the DI virus confers IFN-independent protection against lethal infection [34]. However, the role of type III IFNs in DI-mediated protection has not yet been determined. Additionally, except for physiological characteristics of disease progression, molecular mechanisms underlying DI-mediated protection, especially in the absence of IFN signaling, remain unclear.

In this study, we assessed the efficacy of DI-mediated protection against lethal homologous virus infection in mice deficient in type I and/or type III IFN signaling and investigated the underlying molecular mechanisms. By integrating bulk transcriptomes and microRNA (miRNA) profiles, we identified distinct host signatures associated with DI co-infected mice. These included less intensive expression of inflammatory and immune response genes, and expression of genes involved in multiciliogenesis. Our findings provide mechanistic insight into DI-mediated protection in the absence of type I or III IFNs, and shed light on a novel connection between inflammation, multiciliogenesis, and disease outcome.

## RESULTS

### DI222 virus protects mice from lethal virus infection independent of type I and III IFN signaling

An early study on DI244, a DI virus-derived from a influenza A/Puerto Rico/8/34(H1N1) virus (PR8), [36] indicated type I IFN-independent protection against homologous IAV infection [34]. However, the generalizability of the phenotype in other PR8-derived DIs and the mechanism of protection conferred by DIs in the absence of IFNs remained unknown. To first test the former, we assessed the efficacy *in vivo* of the naturally occurring PR8-derived DI virus, DI222. We previously characterized this virus and demonstrated that DI222 had efficient interfering ability *in vitro* [37]. In brief, we inoculated intranasally C57BL/6 mice—wild-type (WT) or knock-out mouse lines lacking a functional type I IFN receptor (IFNAR^-/-^, KO)—with 1,000 50% tissue culture infectious doses (TCID_50_) of PR8 virus, or a mixture of PR8 and DI viruses at a DI/PR8 ratio of 3,000:1 (maintaining the same dose of PR8 virus in the inoculum as that in the first group).

PR8-infected WT mice suffered from gradual weight loss, with the majority succumbing to infection by day 9 post infection (p.i.) (**Fig. 1a**), while IFNAR^-/-^ mice exhibited more drastic weight loss by day 5 p.i. and all succumbed to infection by day 8 p.i. (**Fig. 1b**). In contrast, all the mice to which DI222 was administered at the moment of PR8 infection (DI222-PR8 co-infection) experienced less severe weight loss (p = 0.0002 in WT mice and p < 0.0001 in IFNAR^-/-^ mice, compared with PR8-only infection) and began re-gaining weight by approximately day 9-10 p.i. (**Fig. 1**). DI244 virus also conferred protection against PR8 infection, although to a lesser degree when compared with the survival rate of the DI222 co-infected mice (p = 0.0398 in IFNAR^-/-^ mice and p = 0.1068 in WT mice, **Fig. 1**). These results demonstrated the efficacy of the DI222 virus in protecting from lethal virus infection, as it led to 100% survival in a type I IFN-independent manner.

**Figure 1.**
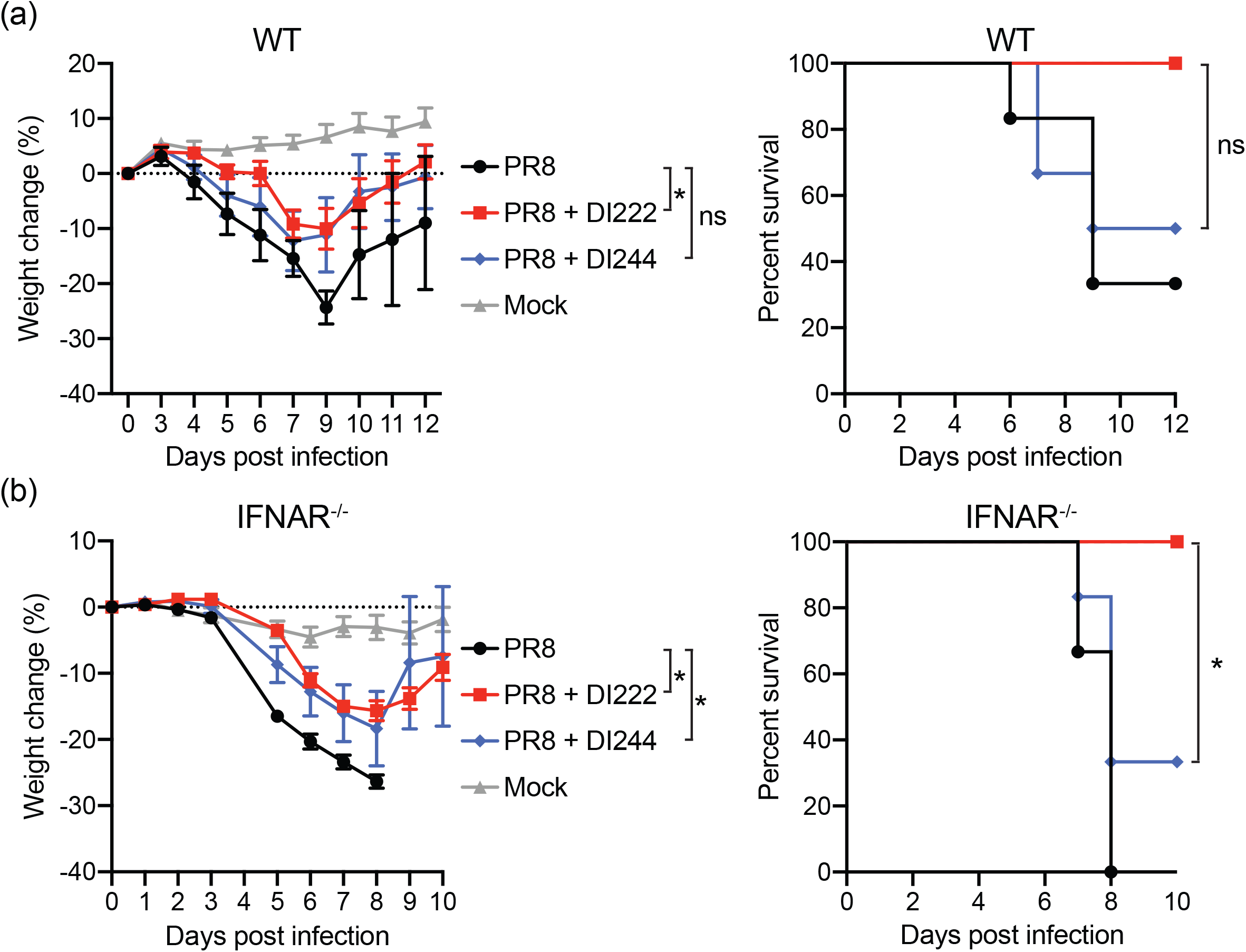
Efficacy of DI222 in protecting mice against lethal PR8 infection independent of type I IFN signaling. Percent weight change (left) and survival (right) were monitored for WT (**a**; n = 5 or 6 per co-infection group) and IFNAR^-/-^ (**b**; n = 6 per co-infection group) mice inoculated with 10^3^ TCID_50_ PR8 virus alone or a mixture of 10^3^ TCID_50_ PR8 and 3 × 10^6^ TCID_50_ DI222 or DI244 viruses. Weight change was analyzed by two-way ANOVA with a mixed-effects model and Šidák correction for multiple comparisons up to day 12 p.i. for WT mice or day 8 p.i. for IFNAR^-/-^ mice. Survival between PR8+DI244 and PR8+DI222 groups was compared using Mantel log-rank analysis and subsequent Bonferroni-Šidák correction for multiple comparisons Data show mean ± standard error of the mean. Asterisk denotes p ≤ 0.05 and “ns” indicates not significant.

Type I and III IFNs are induced following IAV infection and in turn activate antiviral and immunomodulatory responses through a partially overlapping signaling cascade that serves as a foundation for innate immunity in the respiratory epithelium [23]. To determine if both type I and III IFNs are bypassed in DI virus mediated protection in homologous WT IAV infection, we conducted a second study where we included mice lacking the functional type III IFN receptor (IFNLR^-/-^), or both type I and type III receptors (IFNAR^-/-^ IFNLR^-/-^). The inoculation scheme remained the same as that in the first study, and disease severity of infected mice was monitored for 14 days (**Fig. 2a**). PR8 infection generally resulted in rapid weight loss by approximately day 4-5 p.i., accompanied by gradual worsening of symptoms, and substantial mortality in mice deficient in type I or type III IFN signaling (**Fig. 2b and 2c**). However, all the IFNAR^-/-^ IFNLR^-/-^ double knock-out (DKO) mice survived the infection and only manifested mild clinical symptoms, except for a delay in regaining weight compared to PR8-infected IFNLR^-/-^ mice (**Fig. 2d**), suggesting the absence of both type I and III IFN signaling reduces severity of disease in an otherwise lethal infection. In marked contrast, mice from each KO strain co-infected with DI222 virus had even milder symptoms (p < 0.0001 versus PR8-only infection in single KO mice and p = 0.0056 in DKO mice), less weight loss compared with PR8-only infection (p < 0.0001), and 100% survival regardless of the presence or absence of type I and III IFN signaling (**Fig. 2b-d**). Taken together, these results suggest DI-mediated protection against lethal infection with homologous influenza virus is independent of type I and III IFN signaling, while disease progression in PR8-only infection is impacted by the presence or absence of type I and III IFN signaling.

**Figure 2.**
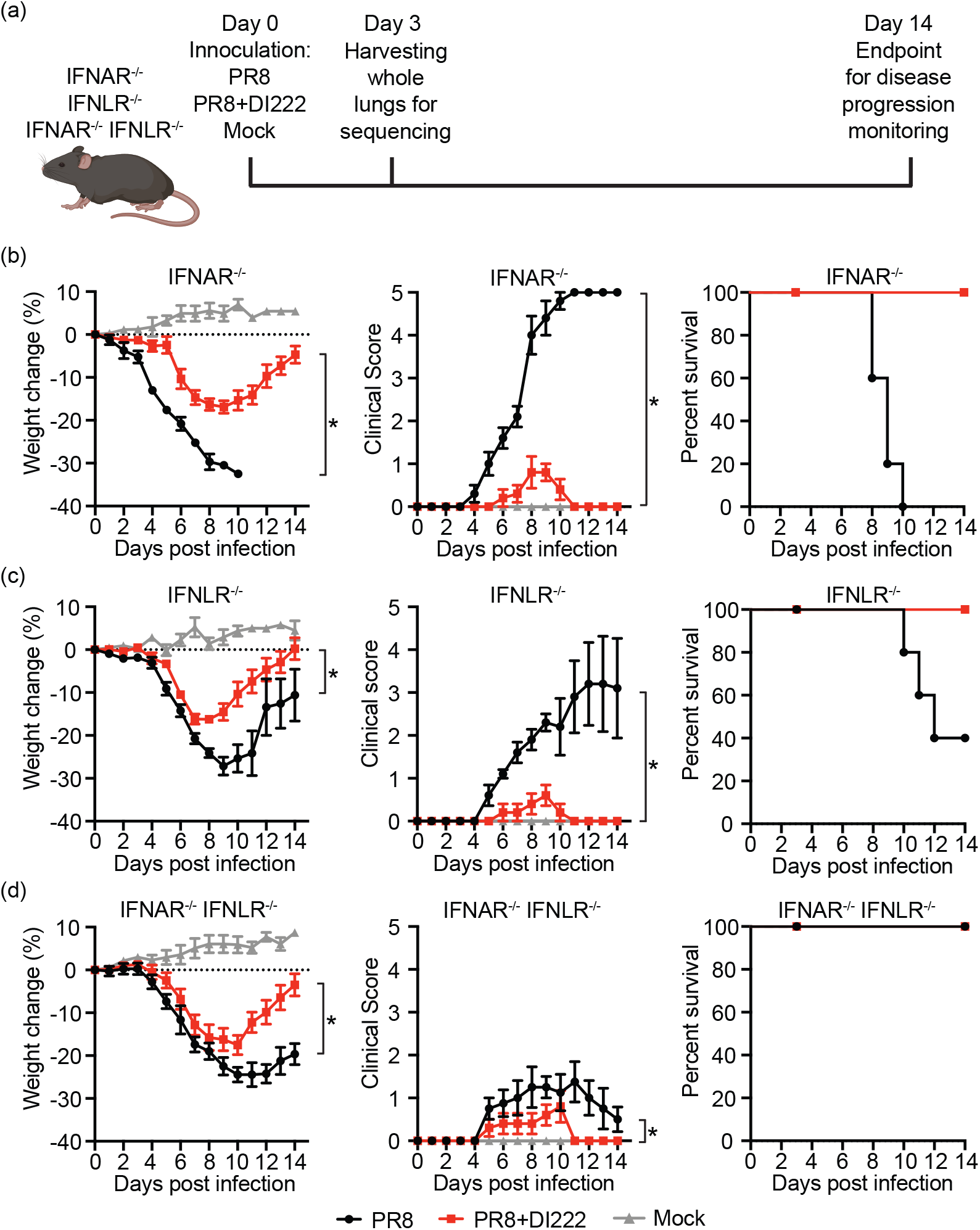
DI222 virus co-infection protects mice from lethal PR8 infection independent of type I and III IFN signaling. (**a**) Experiment setup of DI222 co-infection in different KO mouse strains. Diagram was created in part with BioRender.com. (**b-d**) Percent weight change (left), clinical score (middle), and survival (right) over the course of the infection. IFNAR^-/-^ (**b**; n = 10 per treatment group of which 5 were sacrificed at day 3 p.i. for sequencing), IFNLR^-/-^ (**c**; n = 9 or 11 per treatment group of which 4 or 6 were sacrificed at day 3 p.i.), and IFNAR^-/-^ IFNLR^-/-^ (**d**; n = 7 or 8 per treatment group of which 3 were sacrificed at day 3 p.i.) mice were inoculated with 10^3^ TCID_50_ PR8 virus alone or a mixture of 10^3^ TCID_50_ PR8 and 3 × 10^6^ TCID_50_ DI222 viruses. Another group of mice from each strain (n = 4 in IFNAR^-/-^ and n = 3 for IFNLR^-/-^ and IFNAR^-/-^ IFNLR^-/-^) were mock-infected and sacrificed at day 3 p.i. for sequencing. Disease progression was monitored daily for 14 days. Weight change and clinical score were analyzed by two-way ANOVA with a mixed-effects model and Šidák correction for multiple comparisons up to day 10 p.i. for IFNAR^-/-^ mice or day 14 p.i. for IFNLR^-/-^ and DKO mice. Data show mean ± standard error of the mean. Asterisk denotes p ≤ 0.05 and “ns” indicates not significant.

### Reduced viral transcription, mitigated inflammatory and antiviral responses, and uniquely primed multiciliogenesis in DI co-infected mice

To determine how host expression responses to PR8-only infection and DI222 co-infection could underlie the observed differences in disease severity, we examined mRNA-seq mouse and viral transcriptional profiles in lungs collected at day 3 p.i. We determined lung viral load directly with titration assays and by using virus transcript level estimates as a proxy, calculated as the percentage of viral reads in the total number of reads of a given sample. Viral titers in lung tissue were not statistically different across conditions or mouse strains (**Fig. 3a)**. However, viral transcription levels were generally higher in PR8-only infection compared with that in DI222 co-infection (p < 0.0001 in IFNAR^-/-^ and p = 0.0023 in IFNLR^-/-^ single KO mice, **Fig. 3b**), except in IFNAR^-/-^ IFNLR^-/-^ DKO mice (p = 0.3054, **Fig. 3b**). It is noteworthy that the abundance of DI222 in the viral population in DI222 co-infected mice was maintained at a consistently higher level than that of other defective viral genomes species across strains and at both viral transcriptomic and genomic levels (**Fig. S1**), suggesting persistence of DI222 virus to at least day 3 p.i. We also did a glycomic analysis of the lung tissue to assess the functional potential of the viruses used in the infection, and to determine if glycomic markers of disease severity could be identified, as previously done in the ferret model ([38, 39]). The loss of alpha2,3 sialic acids in lung tissue in the infection groups as compared to the Mock infections confirms viral sialidase activity; there was no statistical difference across the infection groups (**Fig. 3c; Fig. S2**). We also observed higher levels of core 1/3 O-glycans upon infection, particularly in the IFNAR-/-mice, but no statistical difference between the PR8-only and DI222 co-infection groups.

**Figure 3.**
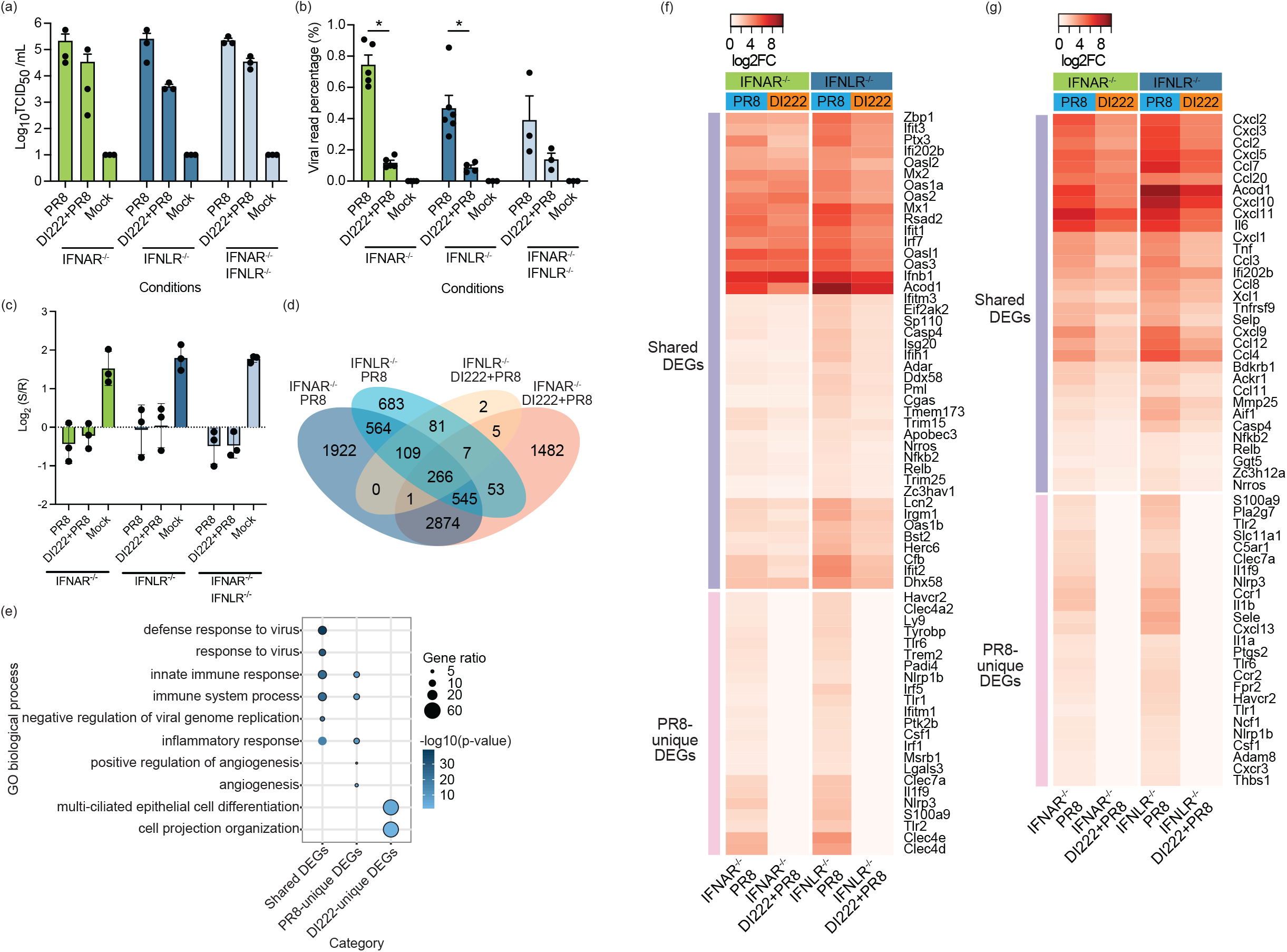
Viral load and host transcriptional response to PR8 infection and DI222 co-infection in mouse lungs at 3 dpi. (**a** and **b**) Viral titers (**a**) and the percentage of viral reads (**b**) in each sample across different mouse strains and virus infection conditions. Each dot represents a sample. Data show means ± standard errors. The significance levels of pairwise comparisons were determined by ordinary one-way ANOVA with Tukey’s multiple-comparison test. Significant differences between virus infection conditions within each mouse strain were denoted by the asterisks (* p ≤ 0.05 and “ns” as not significant). **(c)** Dual-color lectin microarray data for *α* -2,3 sialic acid specific probe SLBRN. Data plotted is median normalized log_2_(sample/reference) data for individual arrays, where the reference is an orthogonally labeled mixture of all samples. (**d)** Venn diagram shows the number of DEGs in PR8-infected and DI222 co-infected IFNAR^-/-^ and IFNLR^-/-^ mice. (**e**) Biological process GO terms associated with up-regulated DEG sets that are commonly induced upon infections or uniquely expressed in response to PR8-infection or DI222 co-infection regardless of the presence of type I or III IFN signaling. The top 5 over-represented GO terms in each DEG set were shown and highlighted by a black outline. The color intensity of each dot indicates the Benjamini p-value and the size corresponds to the percentage of query genes associated with a given GO term in the total number of query genes. (**f**-**g**) Heatmap depicts log2(fold change (FC)) of significantly up-regulated genes involved in the innate immune response (**f**) and inflammatory response (**g**) in each infection group compared with mock-infected group. The genes included have log2FC ≥ 1 in at least one infection group.

To investigate the variation of the host transcriptional landscape under different virus infection conditions and in the presence or absence of type I and III IFN signaling, we first identified differentially expressed genes (DEGs) by comparing virus-infected (both with and without DI) and mock-infected mice. This revealed a larger number of DEGs in PR8-only infection compared with that in the DI222 co-infection, and in IFNAR^-/-^ mice compared with that in IFNLR^-/-^ mice (**Fig. 3d** and **Table S1**). Interestingly, in IFNAR^-/-^ IFNLR^-/-^ DKO mice, only 2 DEGs were detected in PR8 infection and none in the DI222-PR8 co-infection (**Table S1**), suggesting an essential role for type I and III IFNs in initiating host responses to viral infections. To better understand the differences in the host response to each infection condition, regardless of a specific type of IFN, we characterized DEGs that were shared or unique to each condition and detected in both single KO strains. Gene ontology (GO) over-representation analyses of up-regulated DEGs in each set identified shared and distinct host signatures. As expected, IAV infections in general elicited innate immune and inflammatory responses, and defense response to virus, as shown by the shared DEG sets (**Fig. 3e**). PR8-only infection induced additional sets of genes involved in innate immune and inflammatory responses, as well as genes related to angiogenesis and its positive regulation (**Fig. 3e**). In contrast, DI222 co-infection resulted in multi-ciliated epithelial cell differentiation and cell projection organization (**Fig. 3e**), indicated by the expression of the transcription activator Multicilin encoded by *Mcidas* [40], downstream genes involved in centriole biogenesis and assembly, such as *Ccno* [41], *Deup1* [42, 43], and transcription factor *Foxn4* [44, 45]. Interestingly, a more substantial change in the expression level of DEGs involved in the innate immune response (e.g., *Ifit3, Oas1b, Bst2, Ddx58, Rsad2, Isg20*, etc.) and, particularly, inflammation (e.g., *Cxcl3, Cxcl5, Cxcl9, Cxcl10, Ccl2, Ccl3, Ccl4, Ccl7, Ccl12, Acod1* etc.) was primarily observed in PR8-only infection compared with that in DI222 co-infection, as well as in IFNLR^-/-^ mice compared with IFNAR^-/-^ mice infected with PR8 virus alone (**Fig. 3f** and **3g**). The elevated inflammatory response in PR8-only infection can be attributed, at least partially, to monocytes, macrophages, and neutrophils—phagocytes contributing to overt inflammation—as evidenced by a greater enrichment of corresponding cell type gene signatures (**Fig. S3**). These results suggest an increase in the innate immune response and an exuberant inflammatory response are associated with PR8-only infection and the presence of type I IFN signaling, while DI222 co-infection promotes the differentiation of multi-ciliated cells independently of type I or III IFN signaling.

### Integrated miRNA and mRNA co-expression network analysis reveals concurrent miRNA-mRNA signatures associated with DI virus co-infection

Micro RNAs (miRNA) play an important role in regulating gene expression post-transcriptionally and have been implicated in modulating viral replication and host responses during IAV infections (reviewed in [46]). To help better understand host transcriptional response differences between PR8-only infection and DI222 co-infection, we sought to systematically characterize miRNA expression profiles (miRNomes) in the mouse lungs at day 3 p.i. Differential expression analysis of miRNAs (DE-miRNAs) comparing PR8-infected or DI222 co-infected mice with their corresponding mock infection controls indicated changes in the miRNomes similar to that observed in DEG analyses. More DE-miRNAs were identified in PR8-infected IFNAR^-/-^ mice than in DI222 co-infection, and only a few miRNAs were significantly differentially expressed in IFNLR^-/-^ mice (**Fig. 4a** and **Table S2**), while none in DKO mice. Among those infection-responsive miRNAs, three (i.e., miR-223-3p, miR-223-5p, and miR-147-5p) were uniquely up-regulated upon PR8 infection, and one (i.e., miR-449c-5p) in DI222 co-infection, regardless of the deficiency in either type of IFNs (**Fig. 4a** and **Table S2**). These results indicate potential miRNA markers associated with PR8-only infection and DI222 co-infection.

**Figure 4.**
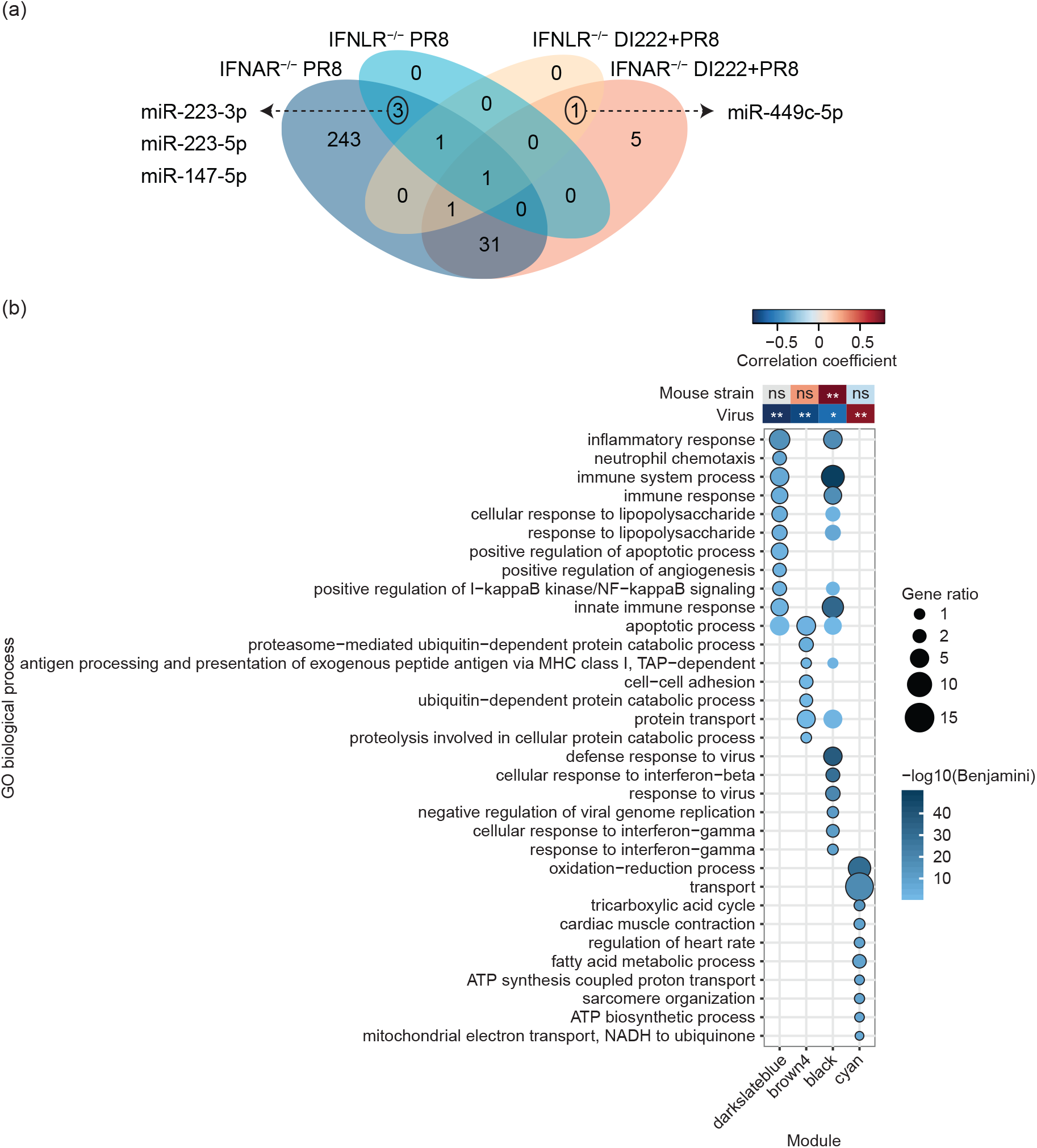
Differentially expressed miRNAs and integrated mRNA-miRNAs signatures associated with virus infection or IFN signaling. (**a**) Venn diagram shows the number of DE-miRNAs in PR8-infected and DI222 co-infected IFNAR^-/-^ and IFNLR^-/-^ mice. (**b**) Heatmap in the upper panel shows the Pearson’s correlation coefficients between modules and traits, including mouse strains (IFNLR^-/-^ versus IFNAR^-/-^) and virus infection conditions (DI222 co-infection versus PR8-only infection), and their significance levels with Benjamini-Hochberg correction denoted by the asterisks (* p ≤ 0.05, ** p ≤ 0.01, *** p ≤ 0.001, **** p ≤ 0.0001, and “ns” as not significant). Bubble plots in the lower panel show over-represented GO terms in a given module. Each GO term is denoted by a dot. The top 10 GO terms in each module were shown and highlighted by a black outline. The color intensity of each dot indicates the Benjamini p-value and the size corresponds to the percentage of query genes associated with a given GO term in the total number of query genes. Module compositions can be found in **Table S3**.

We next aimed to address in a comprehensive manner the relationship between differential expression of mRNAs and miRNAs during IAV infection, and directly evaluate their association with virus infection conditions and type I and III IFN signaling. To do so we constructed an integrated mRNA-miRNA co-expression network using transcriptomic and miRNomic data from IFNAR^-/-^ and IFNLR^-/-^ single KO mice collected at day 3 p.i. Using the WGCNA pipeline [47], we identified modules of co-expressed mRNAs, or both mRNAs and miRNAs, that shared similar expression patterns across conditions. We subsequently correlated these expression patterns with virus infection conditions (DI222-PR8 co-infection versus PR8-only infection) and KO mouse strains (IFNLR^-/-^ versus IFNAR^-/-^) separately to assess the association between modules and infection conditions or IFN signaling. We then performed GO term over-representation analyses with trait-associated modules to determine their biological significance. We identified two modules negatively correlated with virus infection conditions, indicative of gene expression in PR8-only infection: an immune/inflammatory response-related (darkslateblue) module contained genes involved in a broad innate immune response and particularly an inflammatory response, and a module (brown4) contained genes involved in the apoptotic process, protein transport and the ubiquitin-dependent protein catabolic process, as well as antigen processing and MHC class I presentation (**Fig. 4b**). Additionally, another module (black)—comprised of genes primarily involved in the antiviral response, including type I and II IFN responses— negatively correlated with virus infection conditions and positively correlated with mouse strains, representing gene expression in PR8-only infection and in IFNLR^-/-^ mice concurrently (**Fig. 4b**). In contrast, a metabolic-related module (cyan) indicative of gene expression in DI222 co-infection showed over-representation of genes involved in oxidative phosphorylation, muscle contraction, and lipid metabolic process (**Fig. 4b**). It is noteworthy that three DE-miRNAs (miR-223-3p, miR-223-5p, and miR-147-5p) uniquely up-regulated in PR8-only infection in both single KO mouse strains were found in the immune/inflammatory response-related module (darkslateblue), while miR-449c-5p and three multi-ciliated cell-associated DEGs (*Mcidas, Ccno*, and *Foxn4*) uniquely up-regulated in DI222 co-infection were in the metabolic-related module (cyan; **Table S3**). These observations are in line with reports of miR-223 as an immunomodulator and biomarker of some inflammatory diseases (reviewed in [48-50]); and of miR-449, which has a role in promoting multiciliogenesis [51]. Taken together, these results revealed combinatorial expression signatures of mRNAs and miRNAs and further demonstrated a direct link between specific biological processes and different virus infection conditions or the presence of type I or III IFN signaling.

## DISCUSSION

Our data demonstrated type I and III IFN-independent protection conferred by DI virus co-infection, which resulted in reduced viral transcription and promoted multiciliated cell differentiation in mouse lungs at an early stage of the infection. This is in contrast to the inflammatory and innate immune responses marked by intensive expression of inflammatory cytokines and chemokines without DI co-infection. The characteristics of infections with and without DI co-infection are observed in both the transcriptional response and the post-transcriptional regulation by miRNAs, which potentially serve as biomarkers. Taken together, these data provide molecular insights into the mechanism of action of DI virus in conferring protection against lethal homologous virus infection.

DI viruses have been considered appealing candidates as antivirals and vaccine adjuvants given their known functions in pathogenesis (reviewed in [6, 7]). However, gaps in knowledge about the mechanism of action underpinning DI-mediated protection in a clinically relevant system represent one of the major hurdles in developing DI-based prophylactic and therapeutic strategies. Specifically, it remains unclear if the protection *in vivo* arises from direct competition with replication-competent virus, or the strong immunostimulatory ability of a DI virus, including modulation of innate and adaptive immunity as discussed in [6]. Using mice lacking signaling from type I IFNs, type III, or both, we interrogated the role of IFN-modulated innate immune responses in DI-mediated protection against homologous virus infection and characterized the host transcriptional response to DI co-infection in the absence of innate IFN signaling. Our data suggest a link between reduced viral transcription (likely partially due to competition between DI and replication-competent viruses), less severe inflammation, primed muliticiliated cell differentiation, and increased survival in DI co-infection compared with an otherwise lethal infection. It is worth noting that the time point at which the host response was examined in this study only provides a snapshot of pathogenesis during the early stage of infection (i.e. day 3 p.i.), and thus other factors, including, but not limited to, an immediate but transient response triggered even earlier by DI virus independent of type I and III IFNs, and an augmented humoral and cellular response elicited later, as demonstrated in copy-back Sendai virus (SeV) DVGs [52], may also play a role in establishing DI-mediated antiviral protection. Furthermore, given previous reports of reduced efficacy of DI244-mediated protection against infections with influenza B virus [53] and further diminished protection against paramyxovirus infection [34] in the absence of type I IFN signaling, differences in the mechanism of action behind DI-mediated protection against homologous and heterologous infections necessitates additional investigation.

Our integrated characterization of host transcriptional responses at both mRNA and miRNA levels suggests coherent host signatures that differentiate DI co-infection from PR8-only infection. We detected a unique signature of multiciliated cell differentiation in DI co-infection, as evidenced by up-regulation of the transcription regulator Multicilin encoded by *Mcidas* [40, 42, 54] and its downstream targets, including *Ccno* [41], *Deup1* [42, 43], and *Foxn4* [44], involved in centriole biogenesis and assembly during multiciliogenesis. Interestingly, miR-449, which accumulates at a high level in ciliated cells and acts as a key regulator of multiciliogenesis by directly targeting the Delta/Notch pathway that represses Multicilin, in turn de-repressing the adoption of ciliated cell fate [51], is also induced during DI co-infection. These data indicate promoted multiciliated cell differentiation at an early stage of infection with DI co-infection, although the time point is fairly early for epithelial repair, thus a longitudinal study would be needed to determine the relationship between multiciliogenesis and DI treatment, and its implication. Conversely, an augmented inflammatory signature indicated by strong induction of cytokines and chemokines, as well as several miRNA markers were identified in PR8-only infections. Among these miRNAs, miR-223 is known to play a multifaceted role in modulating inflammation, including enhancing granulopoiesis, suppressing neutrophil activation, and directing macrophage polarization and activation towards an anti-inflammatory state (reviewed in [48, 49]). Intriguingly, a higher fold change of miR-223 has been observed in the lungs of mice infected with highly pathogenic influenza virus strains compared with that in less virulent strains [55, 56] and inhibition of miR-223 reduced viral load and mortality [55]. New studies should address the interaction between miR-223 and the inflammatory response during viral infections.

The profiling of innate immune and inflammatory responses across different IFN receptor-deficient mouse strains revealed a stronger response in the absence of type III IFN signaling compared with those lacking type I. This observation is in line with the current paradigm depicting the type III IFN response as being less inflammatory and localized to the site of infection with lower potency in comparison to a systemic type I IFN response [57-59]. Increased morbidity and mortality in mice lacking type I IFN signaling further indicates a critical role for the type I IFN response when the local response is insufficient, highlighting distinct characteristics for these two types of IFNs. Consistent with a previous report of significantly diminished ISG induction in IAV-infected epithelia deficient in receptors of both types of IFNs [23], we observed the response was abolished in mice lacking signaling from both IFNs. However, unlike mice deficient in signaling from both types of IFNs solely in the stromal compartment, who suffered from increased susceptibility to infection, our mice with global deficiency exhibited mitigated morbidity and mortality. This suggests reduced pathogenesis in the systemic absence of type I and III IFN signaling, although the mechanism of pathogen clearance remains unclear. Additional investigations are required to elucidate the tissue-specific IFN response in physiologically-relevant microenvironments during viral infections.

## MATERIALS AND METHODS

### Cells and viruses

Madin-Darby canine kidney (MDCK) cells were maintained in minimum essential medium (MEM) supplemented with 5% FBS. PB2 protein-expressing modified MDCK (AX4/PB2) cells [60] were maintained in MEM supplemented with 5% Newborn Calf Serum (NBCS), 2 μg/ml puromycin and 1 μg/ml blasticidin. Influenza A/Puerto Rico/8/34 (H1N1) virus was propagated in MDCK cells. PR8-derived DI222 virus was generated by reverse genetics as previously described [37] and propagated in AX4/PB2 cells. Influenza viruses were grown in MEM supplemented with 0.15% bovine serum albumin (BSA) fraction V, 1% antibiotic-antimycotic and 1.5 μg/mL TPCK co-treated trypsin. Viral titers were determined by 50% tissue culture infectious dose (TCID_50_) assays in MDCK cells for PR8 virus or AX4/PB2 cells for DI222 virus. The sequences of viral stocks were confirmed by Illumina HiSeq 2500 sequencing.

### Animal studies

In the first study, groups of C57BL/6 WT and IFNAR^-/-^ mice (n = 10), 9-11 weeks old, of both sexes, were anesthetized with inhalational isoflurane and each mouse received (i) 10^3^ TCID_50_ units of PR8 virus, (ii) a mixture of 10^3^ TCID_50_ units of PR8 virus and 3×10^6^ TCID_50_ units of DI244 virus, (iii) a mixture of 10^3^ TCID_50_ units of PR8 virus and 3×10^6^ TCID_50_ units of DI222 virus, or (iv) mock inoculum (PBS and virus growth media), via the i.n. route (in 50 μL PBS). Mice that received mock-infection were used as negative controls and mice that only received PR8 virus were used as positive controls. Mice were monitored daily for weight loss and survival. Mice with a loss of ≥25% of their initial body weight were euthanized. Lungs were collected at day 3 post-infection (p.i.) and stored in RNAlater (Invitrogen) following the manufacturer’s recommendation until use. All animal procedures were performed in accordance with Institutional Animal Care and Use Committee (IACUC) guidelines and have been approved by the IACUC of the Joan & Sanford I. Weill Medical College of Cornell University (Protocol Number 2009–045). All procedures were performed under inhalational isoflurane anesthesia, and every effort was made to minimize suffering.

In the second study with IFNAR^-/-^, IFNLR^-/-^, and DKO mice, mice were lightly anesthetized with 3% inhaled isoflurane and intranasally inoculated with either 10^3^ TCID_50_ units of WT PR8 virus or 10^3^ TCID_50_ units WT PR8 virus plus 10^6.47^ TCID_50_ units DI222 virus in 50 μL PBS. Mock-infected mice received an intranasal inoculation of PBS. Mice were monitored daily for 14 days for weight loss and clinical signs of infection. Clinical signs were scored as follows: 0=alert, active, smooth fur; 1=mildly hunched, slightly ruffled fur but active and alert; 2=hunched and ruffled fur, not active unless stimulated, weight loss apparent; 4=not active when stimulated, rapid breathing, severe weight loss, sunken face, and severely hunched posture, 5=death. Humane endpoints were determined by 30% or greater weight loss and/or clinical signs of 3 or greater. At day 3 p.i, lungs were collected for downstream processing. All animal procedures were performed in accordance with Institutional Animal Care and Use Committee (IACUC) guidelines and have been approved by the IACUC of St. Jude Children’s Hospital (Protocol Number 513).

### Sample preparation and library construction for sequencing

Whole lungs preserved in RNAlater stabilization solution (Invitrogen) were homogenized in QIAzol lysis reagent (Qiagen) and total RNA was extracted using an miRNeasy mini kit (Qiagen) with on-column DNA digestion using RNase-free DNase (Qiagen) according to the manufacturer’s instructions. RNA purity was verified spectrophotometrically and the quality of RNA samples was assessed using the Agilent 2200 TapeStation. RNA-seq libraries were prepared using the NEBNext Ultra II RNA library prep kit for Illumina following poly(A) mRNA enrichment. Libraries were multiplexed and sequenced on an Illumina NextSeq 500 in HighOutput 2×100 bp mode (v2.5). MiRNA sequencing libraries were prepared using QIAseq miRNA library kit for Illumina (Qiagen) and sequenced on the NextSeq 500 in HighOutput 1×75 bp mode (v2.5). Viral genomic RNA (vRNA) in the lung homogenates was amplified in duplicate using multi-segment reverse-transcription PCR (M-RTPCR) [61]. The M-RTPCR product was subjected to Illumina library preparation using the Nextera XT DNA library prep kit (Illumina) and sequenced on the NextSeq 500 in MidOutput 2×150 bp mode (v2.5).

### Lectin microarray analysis

Lung tissue was homogenized and lysates were labeled with NHS-activated AF555 (Thermo Fisher) as described by Pilobello *et al* [62]. Reference sample was prepared by mixing equal amounts of protein from each sample and labeling with NHS-activated AF647 (Thermo Fisher). Printing, hybridizing, and data analysis were performed as described by Pilobello *et al* [62]. Detailed information can be found in Supplemental Tables S4 and S5.

### Computational analyses of RNA-seq, miRNA-seq, and viral genome sequencing data

RNA-seq and virus genome sequencing data was first trimmed with trimmomatic (v0.36) [63] to remove the adaptors and low quality bases. Reads with a minimal length of 36 bases in the trimmed RNA-seq dataset were aligned to the concatenated mouse (GRCm38) and influenza A/Puerto Rico/8/34 (H1N1) reference with STAR (v2.7.3a) [64] with the default parameters, and counted with featureCounts [65] in the Subread package (v1.5.1) [66]. Reads in the virus genome sequencing dataset were aligned to the influenza A/Puerto Rico/8/34 (H1N1) reference with STAR (v2.7.3a) [64]. MiRNA primary quantification was performed in the GeneGlobe Data Analysis Center (Qiagen) for trimming, mapping, and unique molecular indices (UMI) analysis. Although both miRNAs and piRNAs were detected in the miRNA libraries, only miRNAs were retained for the downstream analyses as few piRNAs were differentially expressed. Differential expression analysis was performed by comparing PR8-infected or DI222 and PR8 co-infected mice with the mock-infected mice in a given mouse strain with edgeR (v3.24.3) [67, 68] using the RNA-seq read counts and the miRNA-seq UMI data. Host mRNAs or miRNAs with the adjusted p-value ≤ 0.05 were identified as significantly differentially expressed in each infection condition for a given mouse strain. A complete list of differentially expressed genes (DEGs) and DE-miRNAs can be found in **Table S1-2**. GO term over-representation analysis of the up-regulated DEGs was performed with the online service DAVID (v6.8) [69, 70] by applying the Benjamini p-value cut-off of 0.05. Enrichment score of cell type gene signatures across IFNAR^-/-^ and IFNLR^-/-^ single KO mice were estimated with xCell (v1.1.0) [71].

To identify DVGs in viral populations, split-read analysis was performed to characterize reads that aligned to both ends of a viral segment, with each aligned portion comprised of at least 25 nucleotides in length. Reads with junction coordinates within a 10-nucleotide window were grouped together. Grouped reads with more than 10 read counts were retained. For viral genome sequencing data, reads with junction coordinates that occurred only in one read or in one M-RTPCR replicate were excluded from further analyses. DVG species abundance was calculated as the percentage of split-reads carrying a given deletion junction in the total number of reads aligned to their corresponding segment.

### Integrated miRNA and mRNA co-expressed network analysis

Normalized mRNA and miRNA expression matrices from IFNAR^-/-^ and IFNLR^-/-^ single KO mice were concatenated to construct a weighted gene co-expression network using the WGCNA package (v1.51) in R [47]. A step-by-step network construction and module detection approach was used, which included hierarchical clustering based on a topological overlap matrix (TOM) dissimilarity matrix that was converted from a soft-power-thresholded adjacency matrix (soft-power = 9) and subsequent hybrid dynamic tree-cutting (deepSplit = 3) with a minimal module size constraint of 20. Highly similar modules were further merged at a cut height of 0.25. The module eigengene (ME) defined as the first principal component of a module was used to detect trait-associated modules by establishing Pearson’s correlation with binary traits, including mouse strain (IFNAR^-/-^: 0; IFNLR^-/-^: 1) and virus (PR8: 0; DI222 + PR8: 1). GO term over-representation analysis of the genes in trait-associated modules was performed with the online service DAVID (v6.8) as described above. A complete list of module compositions can be found in **Table S3**.

### STATISTICAL ANALYSIS

Weight change related statistical analyses were performed as indicated in figure legends using Prism 9 (GraphPad Software). Pairwise comparisons of survival between designated groups were conducted with Mantel log-rank analysis and subsequent Bonferroni-Šidák correction for multiple comparisons. The statistical significance of the difference in the viral titer and relative abundance of viral transcripts between two groups of mice was determined by ordinary one-way ANOVA with Tukey’s multiple-comparison test. A p-value of ≤ 0.05 was considered statistically significant.

## CODE AVAILABILITY

The code used to generate all the results is available on Github (https://github.com/GhedinLab/DI_IFNR-KO-Mice).

## DATA AVAILABILITY

Sequencing data that support the findings of this study have been deposited in the Gene Expression Omnibus (GEO) repository under accession GSE18122. Lectin microarray data can be found at doi: 10.7303/syn26242399.

## ACKNOWLEDGEMENTS

We thank members of the Ghedin laboratory for feedback and discussion. We thank Olivia Micci-Smith, Hana Husic, Nicole Adamski, and Mohammed Khalfan of the Genomics Core Facility at the Center for Genomics and Systems Biology, New York University. We thank Dr. Yoshihiro Kawaoka for providing the AX4/PB2 cells. This work was supported in part by NIAID/NIH R01AI140766 (E.G., S.S.C), CIVIC HHS-NIH-NIAID-BAA2018 (S.S.C.), CEIRR 75N93021C00016 and CEIRS HHSN27220140006C (S.S.C), R21AI124141 (M.S.), the Canada Excellence Research Chairs Program (L.K.M.). and the Intramural Research Program of the NIAID/NIH (E.G.). This work was also supported in part through the NYU IT High Performance Computing resources, services, and staff expertise.

## AUTHOR CONTRIBUTIONS

C.W. and E.G. conceived and designed the study, with input from S.S.C. and M.S. The experiments were performed by R.H., M.S, JY, LM, and NMT. Sequencing libraries preparations were carried out by C.W. The resulting sequencing data was analyzed by C.W. and analysis of other types of data were done by R.H. and C.W. The manuscript was written by C.W. and E.G., with input from all authors. All the authors viewed and approved the manuscript. The study was supervised by E.G. and S.S.C. Funding acquisition was done by E.G., S.S.C. and M.S.

## SUPPLEMENTARY FIGURE LEGENDS

**Figure S1.**
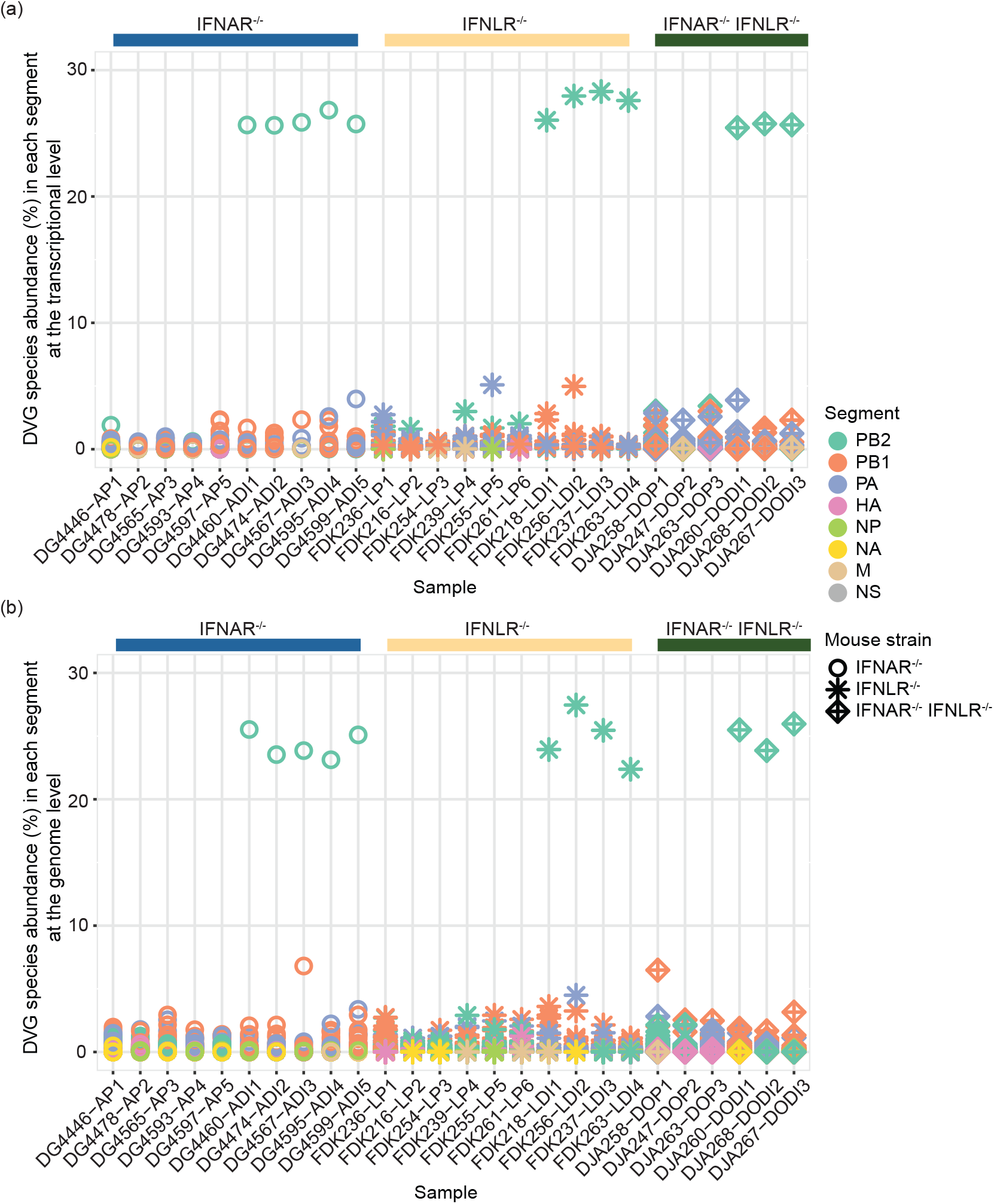
Distribution of DVG abundance in each segment across different virus infection conditions and mouse strains. Dots depict the percentage of split reads derived from a given DVG species in its corresponding segment across samples at viral transcriptomic (**a**) and genomic (**b**) levels. The color indicates viral segments and the shape corresponds to mouse strains.

**Figure S2.**
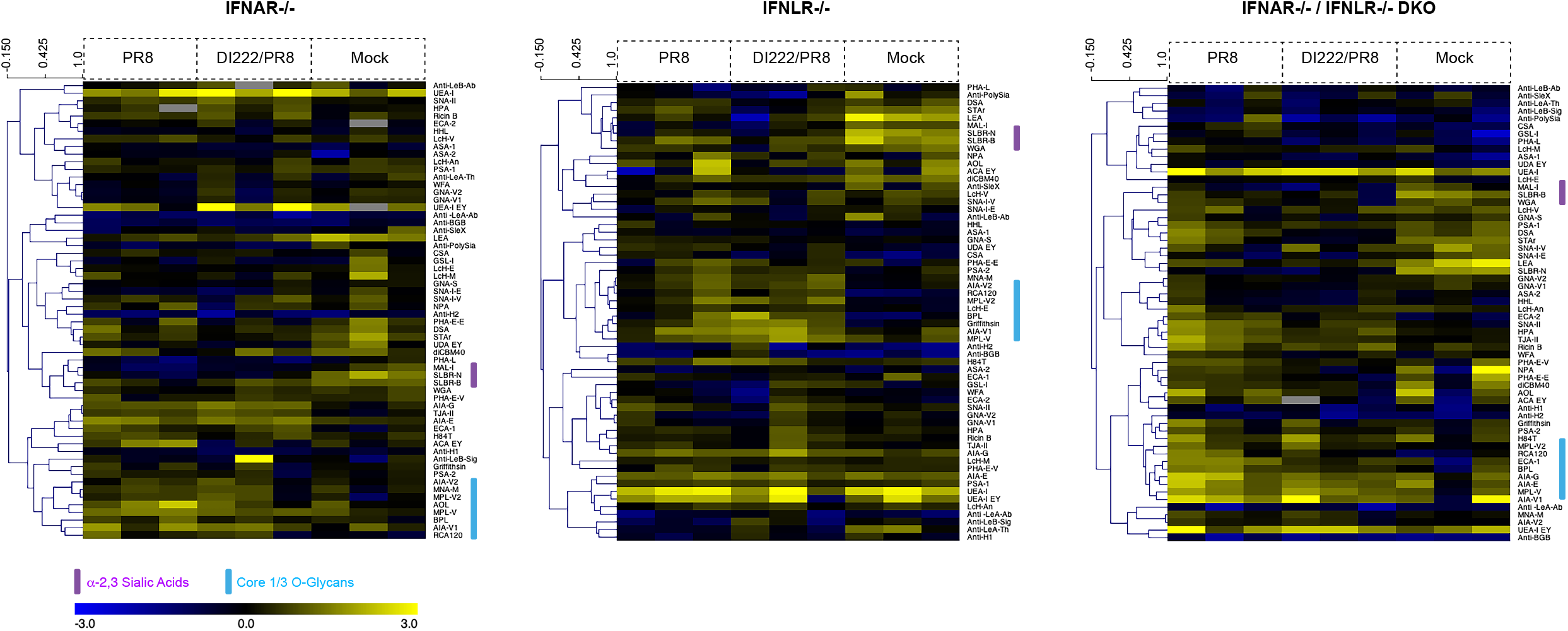
Heat maps of lectin microarray analysis of mouse lungs across different virus infection conditions and mouse strains. Heatmaps were generated for each mouse strain (IFNAR-/-, IFNLR-/-, IFNAR-/-/IFNLR-/-). Median normalized log_2_ ratios (Sample (S)/Reference(R)) of mouse lung samples were ordered by infection (PR8-pink, DI22/PR8-green, Mock-blue). A common mixed biological reference consisting of samples from all conditions and mouse strains was used. Yellow, log_2_(S) > log_2_(R); blue, log_2_(R) > log_2_(S). Lectins binding *α*-2,3-sialosides (magenta) and core 1/3 O-glycans (light blue) are indicated.

**Figure S3.**
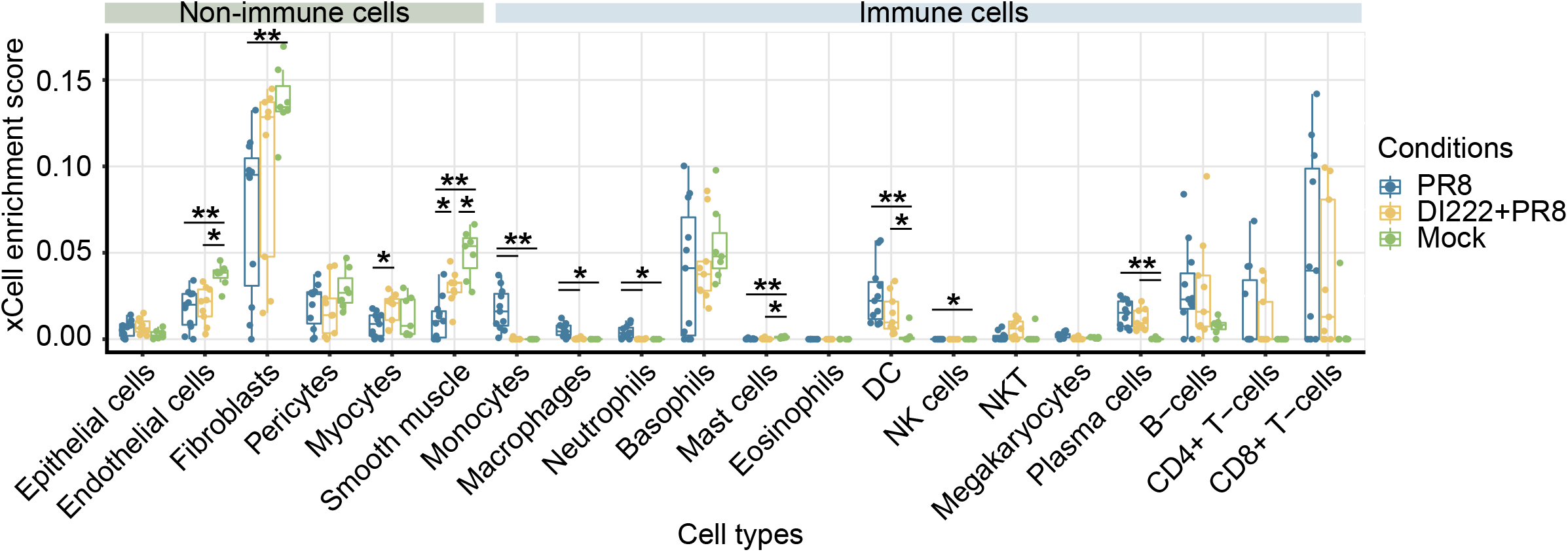
Enrichment of cell type gene signatures across different virus infection conditions. Enrichment scores of a given cell type gene signature in the IFNAR^-/-^ and IFNLR^-/-^ single KO mice grouped by virus infection conditions across a wide variety of cell types were estimated using xCell. Dots denote the enrichment score of a sample. Boxes show the inter-quartile range (IQR) with the median in the center and whiskers are minimum to maximum within 1.5 x IQR from the first and third quartiles. The color of dots and boxes indicates virus infection conditions. The significance levels of pairwise comparisons between groups (i.e., infection conditions) for a given cell type are determined by two-tailed Wilcoxon rank sum test with Benjamini-Hochberg correction were denoted by the asterisks (* p ≤ 0.05 and ** p ≤ 0.01).

## SUPPLEMENTARY TABLES

**Table S1. Differentially expressed genes in each mouse strains under different infection conditions**.

**Table S2. Differentially expressed miRNAs in each mouse strains under different infection conditions**.

**Table S3. Module compositions of mRNA-miRNA co-expression network**.

**Table S4. MIRAGE Lectin Microarray Data**.

**Table S5. Lectin microarray print list**.

## REFERENCES

1. Henle, W. and G. Henle, Interference of inactive virus with the propagation of virus of influenza A. Science, 1943.98(2534): p. 87-89.

2. von Magnus, P., Incomplete Forms of Influenza Virus, in Advances in Virus Research Volume 2. 1954. p. 59–79.

3. Huang, A.S. and D. Baltimore, Defective viral particles and viral disease processes. Nature, 1970. 226: p. 325–327.

4. Lazzarini, R.A., J.D. Keene, and M. Schubert, The origins of defective interfering particles of the negative-strand RNA viruses. Cell, 1981. 26(2): p. 145-154.

5. Genoyer, E. and C.B. López, The Impact of Defective Viruses on Infection and Immunity. Annual Review of Virology, 2019. 6: p. 547–566.

6. Vignuzzi, M. and C.B. López, Defective viral genomes are key drivers of the virus–host interaction. Nature Microbiology, 2019. 4(7): p. 1075-1087.

7. Dimmock, N.J. and A.J. Easton, Defective interfering influenza virus RNAs: time to reevaluate their clinical potential as broad-spectrum antivirals? Journal of Virology, 2014. 88(10): p. 5217-27.

8. Marriott, A.C. and N.J. Dimmock, Defective interfering viruses and their potential as antiviral agents. Reviews in Medical Virology, 2010. 20(1): p. 51-62.

9. Janda, M.J., et al., Diversity and generation of defective interfering influenza virus particles. Virology, 1979. 95(1): p. 48-58.

10. Cairns, H.J.F. and M. Edney, Quantitative aspects of influenza virus multiplication. The Journal of Immunology, 1952. 69(2): p. 155-160.

11. Bellett, A.J.D. and P.D. Cooper, Some properties of the transmissible interfering component of vesicular stomatitis virus preparations. The Journal of General Microbiology, 1959. 21: p. 498–509.

12. Sokol, F., A.R. Neurath, and J. Vilcek, Formation of incomplete sendai virus in embryonated eggs. Acta Virologica, 1964. 8: p. 59–67.

13. Saira, K., et al., Sequence analysis of in vivo defective interfering-like RNA of influenza A H1N1 pandemic virus. Journal of Virology, 2013. 87(14): p. 8064-74.

14. Vasilijevic, J., et al., Reduced accumulation of defective viral genomes contributes to severe outcome in influenza virus infected patients. PLos Pathogens, 2017. 13(10): p. e1006650.

15. Sun, Y., et al., Immunostimulatory defective viral genomes from respiratory syncytial virus promote a strong innate antiviral response during infection in mice and humans. PloS Pathogens, 2015. 11(9): p. e1005122.

16. Rabinowitz, S.G. and J. Huprikar, The influence of defective-interfering particles of the PR-8 strain of influenza A virus on the pathogenesis of pulmonary infection in mice. The Journal of Infectious Diseases, 1979. 140(3): p. 305-315.

17. Gamboa, E.T., et al., Murine influenza virus encephalomyelitis. III. Effect of defective interfering virus particles. Acta Neuropathol, 1976. 34(2): p. 157-169.

18. Tapia, K., et al., Defective viral genomes arising in vivo provide critical danger signals for the triggering of lung antiviral immunity. PLos Pathogens, 2013. 9(10): p. e1003703.

19. Manzoni, T.B. and C.B. López, Defective (interfering) viral genomes re-explored: impact on antiviral immunity and virus persistence. Future Virology, 2018. 13(7): p. 493-504.

20. Baum, A., R. Sachidanandam, and A. García-Sastre, Preference of RIG-I for short viral RNA molecules in infected cells revealed by next-generation sequencing. PNAS, 2010. 107(37): p. 16303-16308.

21. Strähle, L., et al., Activation of the beta interferon promoter by unnatural sendai virus infection requires RIG-I and is inhibited by viral C proteins. Journal of Virology, 2007. 81(22): p. 12227-12237.

22. Xu, J., et al., Identification of a natural viral RNA motif that optimizes sensing of viral RNA by RIG-I. mBio, 2015. 6(5): p. e01265-15.

23. Crotta, S., et al., Type I and type III interferons drive redundant amplification loops to induce a transcriptional signature in influenza-infected airway epithelia. PLoS Pathogens, 2013. 9(11): p. e1003773.

24. Iwasaki, A. and P.S. Pillai, Innate immunity to influenza virus infection. Nature Reviews Immunology, 2014. 14(5): p. 315-28.

25. Schoggins, J.W. and C.M. Rice, Interferon-stimulated genes and their antiviral effector functions. Current Opinion in Virology, 2011. 1(6): p. 519-25.

26. Schneider, W.M., M.D. Chevillotte, and C.M. Rice, Interferon-stimulated genes: a complex web of host defenses. Annual Review of Immunology, 2014. 32: p. 513–45.

27. Schoggins, J.W., Interferon-stimulated genes: What do they all do? Annual Review of Virology, 2019. 6: p. 567–584.

28. Jewell, N.A., et al., Lambda interferon is the predominant interferon induced by influenza A virus infection in vivo. Journal of Virology, 2010. 84(21): p. 11515–11522.

29. Okabayashi, T., et al., Type-III interferon, not type-I, is the predominant interferon induced by respiratory viruses in nasal epithelial cells. Virus Research, 2011. 160(1-2): p. 360-366.

30. Fox, J.M., et al., Interferon lambda upregulates IDO1 expression in respiratory epithelial cells after influenza virus infection. Journal of Interferon & Cytokine Research, 2015. 35(7): p. 554-562.

31. Galani, I.E., et al., Interferon-λ mediates non-redundant front-line antiviral protection against influenza virus infection without compromising host fitness. Immunity, 2017. 46(5): p. 875-890.e6.

32. Espinosa, V., et al., Type III interferon is a critical regulator of innate antifungal immunity. Science Immunology, 2017. 2(16): p. eaan5357.

33. Klinkhammer, J., et al., IFN-λ prevents influenza virus spread from the upper airways to the lungs and limits virus transmission. eLife, 2018. 7: p. e33354.

34. Easton, A.J., et al., A novel broad-spectrum treatment for respiratory virus infections: influenza-based defective interfering virus provides protection against pneumovirus infection in vivo. Vaccine, 2011. 29(15): p. 2777-84.

35. Scott, P.D., et al., Defective interfering influenza virus confers only short-lived protection against influenza virus disease: Evidence for a role for adaptive immunity in DI virus-mediated protection in vivo. Vaccine, 2011. 29: p. 6584–6591.

36. Dimmock, N.J., et al., Influenza virus protecting RNA: an effective prophylactic and therapeutic antiviral. Journal of Virology, 2008. 82(17): p. 8570-8578.

37. Wang, C., et al., Cell-to-cell variation in defective virus expression and effects on host responses during influenza virus infection. mBio, 2020. 11: p. e02880–19.

38. Heindel, D.W., et al., Glycomic analysis of host response reveals high mannose as a key mediator of influenza severity. PNAS, 2020. 117(43): p. 26926-26935.

39. Chen, S., et al., Age-Dependent Glycomic Response to the 2009 Pandemic H1N1 Influenza Virus and Its Association with Disease Severity. Journal of Proteome Research, 2020. 19(11): p. 4486-4495.

40. Stubbs, J.L., et al., Multicilin promotes centriole assembly and ciliogenesis during multiciliate cell differentiation. Nature Cell Biology, 2012. 14: p. 140–147.

41. Boon, M., et al., MCIDAS mutations result in a mucociliary clearance disorder with reduced generation of multiple motile cilia. Nature Communications, 2014. 5: p. 4418.

42. Ma, L., et al., Multicilin drives centriole biogenesis via E2f proteins. Genes & Developments, 2014. 28(13): p. 1461-1471.

43. Zhao, H., et al., The Cep63 paralogue Deup1 enables massive de novo centriole biogenesis for vertebrate multiciliogenesis. Nature Cell Biology, 2013. 15(12): p. 1434-1444.

44. Campell, E.P., I.K. Quigley, and C. Kintner, Foxn4 promotes gene expression required for the formation of multiple motile cilia. Development, 2016. 143(24): p. 4654-4664.

45. Li, S. and M. XIang, Foxn4 influences alveologenesis during lung development. Developmental Dynamics, 2011. 240(6): p. 1512-1517.

46. Keshavarz, M., et al., miRNA-based strategy for modulation of influenza A virus infection. Epigenomics, 2018. 10(6): p. 829-844.

47. Langfelder, P. and S. Horvath, WGCNA: an R package for weighted correlation network analysis. BMC Bioinformatics, 2008. 9: p. 559.

48. Haneklaus, M., et al., miR-223: infection, inflammation and cancer. Journal of Internal Medicine, 2013. 274(3): p. 215-26.

49. Yuan, X., et al., MicroRNA miR-223 as regulator of innate immunity. Journal of Leukocyte Biology, 2018. 104(3): p. 515-524.

50. Dunaeva, M., et al., Circulating serum miR-223-3p and miR-16-5p as possible biomarkers of early rheumatoid arthritis. Clinical and Experimental Immunology, 2018. 193(3): p. 376-385.

51. Marcet, B., et al., Control of vertebrate multiciliogenesis by miR-449 through direct repression of the Delta/Notch pathway. Nature Cell Biology, 2011. 13(6): p. 693-9.

52. Mercado-López, X., et al., Highly immunostimulatory RNA derived from a Sendai virus defective viral genome. Vaccine, 2013. 31(48): p. 5713-5721.

53. Scott, P.D., et al., Defective interfering influenza A virus protects in vivo against disease caused by a heterologous influenza B virus. Journal of General Virology, 2011. 92: p. 2122–2132.

54. Kim, S., et al., Multicilin and activated E2f4 induce multiciliated cell differentiation in primary fibroblasts. Scientific Reports, 2018. 8(1): p. 12369.

55. Choi, E.-J., et al., Differential microRNA expression following infection with a mouse-adapted, highly virulent avian H5N2 virus. BMC Microbiology, 2014. 14: p. 252.

56. Vela, E.M., et al., MicroRNA expression in mice infected with seasonal H1N1, swine H1N1 or highly pathogenic H5N1. Journal of Medical Microbiology, 2014. 63: p. 1131–1142.

57. Wack, A., E. Terczynska-Dyla, and R. Hartmann, Guarding the frontiers: the biology of type III interferons. Nature Immunology, 2015. 16(8): p. 802-9.

58. Kotenko, S.V. and J.E. Durbin, Contribution of type III interferons to antiviral immunity: location, location, location. Journal of Biological Chemistry, 2017. 292(18): p. 7295-7303.

59. Wells, A.I. and C.B. Coyne, Type III interferons in antiviral defenses at barrier surfaces. Trends in Immunology, 2018. 39(10): p. 848-858.

60. Ozawa, M., et al., Replication-incompetent influenza A viruses that stably express a foreign gene. Journal of General Virology, 2011. 92: p. 2879–2888.

61. Zhou, B., et al., Single-reaction genomic amplification accelerates sequencing and vaccine production for classical and swine origin human influenza A viruses. Journal of Virology, 2009. 83(19): p. 10309-10313.

62. Pilobello, K.T., et al., Advances in lectin microarray technology: optimized protocols for piezoelectric print conditions. Current Protocols in Chemical Biology, 2013. 5(1): p. 1-23.

63. Bolger, A., M. Lohse, and B. Usadel, Trimmomatic: a flexible trimmer for Illumina sequence data. Bioinformatics, 2014. 30(15): p. 2114-20.

64. Dobin, A., et al., STAR: ultrafast universal RNA-seq aligner. Bioinformatics, 2013. 29(1): p. 15-21.

65. Liao, Y., G. Smyth, and W. Shi, featureCounts: an efficient general purpose program for assigning sequence reads to genomic features. Bioinformatics, 2014. 30(7): p. 923-30.

66. Liao, Y., G. Smyth, and W. Shi, The Subread aligner: fast, accurate and scalable read mapping by seed-and-vote. Nucleic Acids Research, 2013. 41(10): p. e108.

67. Robinson, M., D. McCarthy, and G. Smyth, edgeR: a Bioconductor package for differential expression analysis of digital gene expression data. Bioinformatics, 2010. 26(1): p. 139-140.

68. McCarthy, D., Y. Chen, and G. Smyth, Differential expression analysis of multifactor RNA-seq experiments with respect to biological variation. Nucleic Acids Research, 2012. 40(10): p. 4288-4297.

69. Huang, D.W., B.T. Sherman, and R.A. Lempicki, Systematic and integrative analysis of large gene lists using DAVID Bioinformatics Resources. Nature Protocol, 2009. 4(1): p. 44-57.

70. Huang, D.W., B.T. Sherman, and R.A. Lempicki, Bioinformatics enrichment tools: paths toward the comprehensive functional analysis of large gene lists. Nucleic Acids Research, 2009. 37(1): p. 1-13.

71. Aran, D., Z. Hu, and A.J. Butte, xCell: digitally portraying the tissue cellular heterogeneity landscape. Genome Biology, 2017. 18: p. 220.

